# 25-hydroxycholesterol promotes brain endothelial dysfunction by remodelling cholesterol metabolism

**DOI:** 10.1101/2024.05.10.590792

**Authors:** Victor S. Tapia, Sarah E. Withers, Ran Zhou, Abigail Bennington, Frances Hedley, Adam El Khouja, Nadim Luka, Marco Massimo, Siobhan Crilly, Katherine R. Long, Catherine B. Lawrence, Paul R. Kasher

**Author notes:** **Corresponding author:** Paul Kasher, address: AV Hill building, Oxford Road, Manchester M13 9PT, United Kingdom.

## Abstract

The antiviral enzyme cholesterol 25-hydroxylase (CH25H) and its metabolite 25-hydroxycholesterol (25HC), which modulates cholesterol metabolism during infection, have been previously associated with vascular pathology. Viral infections have been linked to risk of intracerebral haemorrhage (ICH) but the molecular mechanisms leading to brain vessel rupture via antiviral responses remain unknown. We hypothesised that the CH25H/25HC pathway may impact neuroendothelial integrity in the context of infection-associated ICH. Here, using a SARS-CoV-2-spike-induced zebrafish ICH model and foetal human SARS-CoV-2-associated cortical tissue containing microbleeds, we identified an upregulation of CH25H in infection-associated cerebral haemorrhage. Using zebrafish ICH models and human brain endothelial cells, we asked whether 25HC may promote neurovascular dysfunction by modulating cholesterol metabolism. We found that 25HC and pharmacological inhibition of *HMGCR* by atorvastatin interacted to exacerbate brain bleeding in zebrafish larvae and *in vitro* brain endothelial dysfunction. *In vitro* 25HC-induced dysfunction was also rescued by cholesterol supplementation. These results demonstrate that the antiviral factor 25HC can dysregulate brain endothelial function by remodelling cholesterol metabolism. We propose that the CH25H/25HC pathway represents an important component in the pathophysiology of brain vessel dysfunction associated with infection and cholesterol dysregulation in the context of ICH.

**Summary Statement:** The antiviral metabolite 25-hydroxycholesterol dysregulates brain endothelial function by remodelling cholesterol metabolism, thereby providing a mechanistic link between viral infection and brain endothelial dysfunction in conditions such as intracerebral haemorrhage.

## Introduction

Intracerebral haemorrhage (ICH) is a type of stroke caused by the rupture of brain vessels and subsequent bleeding within the brain parenchyma. While several ICH risk factors have been described (1, 2), how they lead to cerebrovascular dysfunction is not completely understood. SARS-CoV-2, herpes zoster, herpes simplex, hepatitis C and dengue infections have been associated with the incidence of ICH (3-7), while flu-like symptoms have been proposed to directly trigger ICH (8). Alongside adult-onset ICH, viral infections may also induce brain haemorrhages during development. Foetal brain haemorrhages have been associated with SARS-CoV-2 infection (9) and maternal immune activation (10). Insights into how antiviral responses lead to brain vessel dysfunction will improve the understanding of haemorrhagic stroke pathophysiology.

The effects of antiviral signalling in the developing brain can also be observed in monogenic conditions such as the type I interferonopathies. Early post-natal brain haemorrhages have been described for Aicardi-Goutières syndrome 5 (AGS5) (11, 12) caused by recessive mutations in the viral restriction factor SAMHD1. We have previously modelled the cerebrovascular phenotype of AGS5 in zebrafish larvae (13, 14), reporting a novel link between type I interferon (IFN) signalling, cholesterol dysregulation and susceptibility to brain haemorrhages (13). Although others have previously highlighted a link between IFN signalling and modulation of cholesterol metabolism (15), this has not been previously associated with cerebrovascular defects.

A mechanistic link between infection and remodelling of cholesterol metabolism has been reported for the IFN-stimulated enzyme cholesterol 25-hydroxylase (CH25H) and its metabolite 25-hydroxycholesterol (25HC) (16). The antiviral mechanisms of 25HC have been well described, involving the inhibition of cholesterol synthesis and depletion of plasma membrane cholesterol (16-18). Besides a role in immune defence, recent studies have shown that the CH25H/25HC pathway has a detrimental role in vascular function in certain pathologies, such as atherosclerosis, lung inflammation and experimental autoimmune encephalomyelitis (EAE) (19-21). Whether CH25H/25HC has a role in cerebrovascular dysfunction associated with ICH remains unknown.

In this study, we hypothesised that cholesterol remodelling induced by 25HC could induce cerebrovascular dysfunction. First, we characterised an upregulation of CH25H in SARS-CoV-2-spike injected zebrafish larvae and human SARS-CoV-2-associated developmental brain haemorrhages. Then we evaluated the effects of 25HC on zebrafish ICH models and in the human brain EC line hCMEC/D3. We found an interaction between 25HC and pharmacological inhibition of *HMGCR* by statins, which increased the severity of brain bleeding in zebrafish larvae and *in vitro* endothelial dysfunction. 25HC effects were also dependent on cholesterol availability, as cholesterol supplementation rescued the 25HC-induced dysfunction in HCMEC/D3 cells. We propose that the CH25H/25HC pathway may be a relevant factor in infection-triggered brain EC dysfunction.

## Results

### CH25H is expressed in infection-associated developmental intracerebral haemorrhage

To ultimately assess the role of the CH25H/25HC pathway in cerebrovascular dysfunction, we wanted to evaluate the expression of CH25H in infection-associated ICH. For this, we took advantage of a SARS-CoV-2 neuroinflammatory model in zebrafish larvae. Although SARS-CoV-2 is not able to replicate in zebrafish (22), injection of the SARS-CoV-2 spike protein (spike) into the hindbrain of zebrafish larvae induces an inflammatory response and brain haemorrhages (23, 24). To confirm these findings, we used a zebrafish double transgenic reporter for endothelial cells and erythrocytes (Tg(fli1:EGFP)/Tg(gata1a:DsRed)) to evaluate spike-induced brain haemorrhages *in vivo* (Fig. 1A).

**Figure 1.**
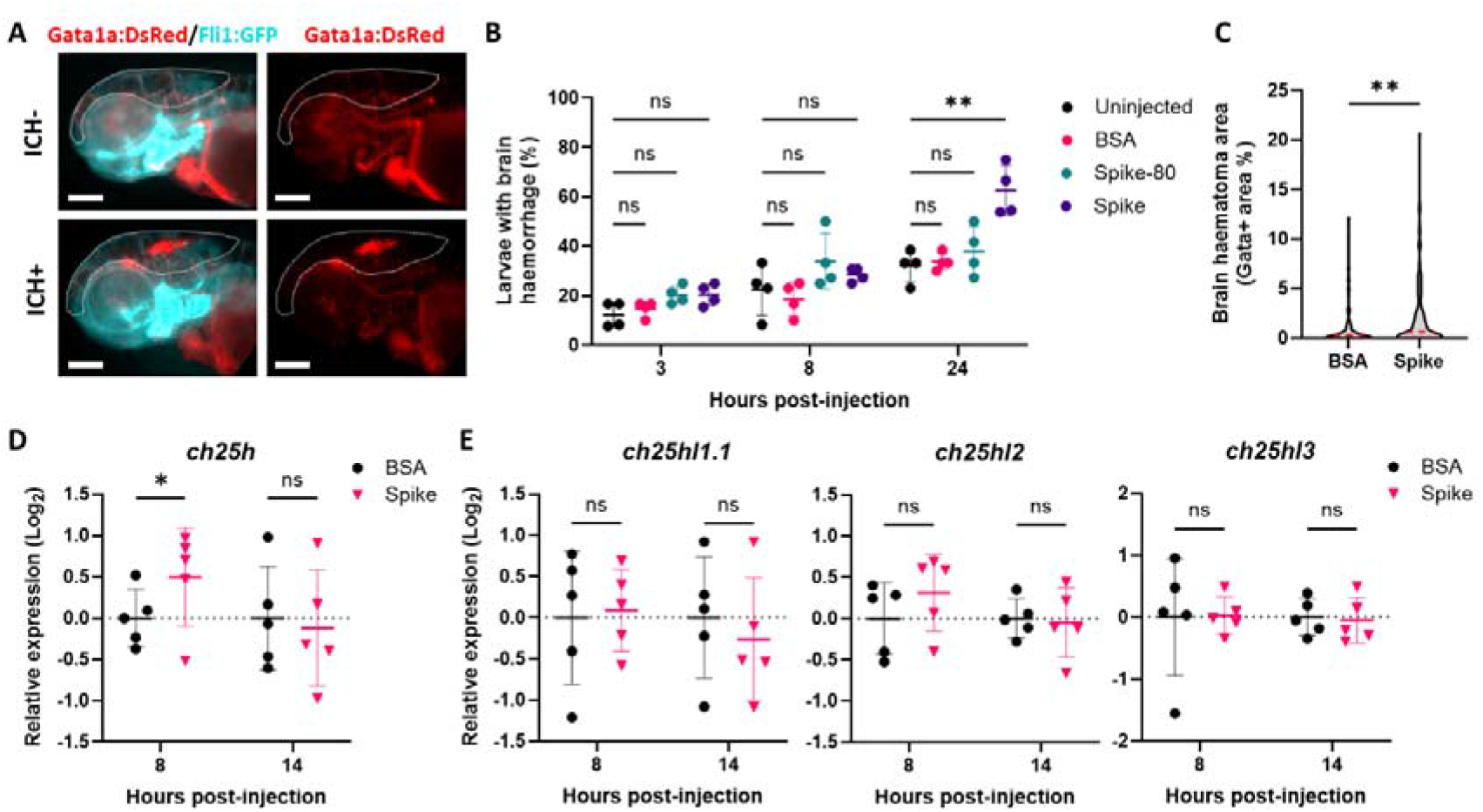
SARS-CoV-2 spike protein induces ch25h upregulation prior to spontaneous brain bleeding in zebrafish larvae. **A** Representative images of control (ICH-) and haemorrhagic (ICH+) *Tg(fli1:EGFP)/Tg(gata1a:DsRed)* 3 days post-fertilization (dpf) larvae, 24 hours after SARS-CoV-2 spike protein (spike) injection into the hindbrain (0.25 mg/ml, 2 nl). Red indicates erythrocytes (gata1a positive) and cyan endothelial cells (fli1+ positive). Dotted lines indicate brain area. Scale bars 250 µm. **B** Time course of ICH+ frequency in *Tg(fli:EGFP)/Tg(gata1a:DsRed)* larvae in uninjected, injected with BSA control, injected with pre-heated spike at 80 °C for 30 min (spike-80) or injected with spike conditions (n=4, 11-14 embryos per experiment). **C** Haematoma size (gata positive area in brain region) in larvae 24 h post-injection of BSA or spike (n=146-147 embryos, 3 independent experiments). **D-E** Gene expression was analysed in larval heads 8 and 14 hours after BSA or spike injections, for *ch25h* (**D**), *ch25hl1*.*1, ch25hl2* and *ch25hl3* (**E**). Data are mean ± SD (**B, C, E**) or median ± IQR (***C***). *, p < 0.05; **, p < 0.01; ns, non-significant, determined by repeated measures ANOVA with Dunnet’s post-hoc test compared to uninjected (**B**), Mann-Whitney test (**C**), or randomized block two-way ANOVA with Sidak’s post hoc analysis compared to BSA (**D, E**).

Spike injections were compared to non-injected, BSA or denatured (80 °C, 30 min) spike protein (spike-80) injected control groups. Spike injection did not change brain haemorrhage frequency at 3 or 8 hours post-injection but induced a significant increase in frequency at 24 hours (Fig. 1B). Furthermore, a significant increase in brain haematoma area was also observed at 24 hours post-injection in spike-injected larvae compared to BSA controls (Fig. 1C). We then evaluated the expression of the five *CH25H* homolog genes described in zebrafish (25). *Ch25h* was transiently upregulated by spike injection before the increase in brain haemorrhage frequency, as observed by a significant upregulation at 8 hours post-injection and return to basal levels at 14 hours (Fig. 1D). In contrast, spike injection had no effect on the other zebrafish homolog genes *ch25hl1*.*1, ch25hl2* and *ch25hl3* (Fig. 1E) and *ch25hl1*.*2* was not detected (not shown). These findings corroborate previous research indicating that zebrafish *ch25h* is upregulated in response to viral challenge, whereas the other *ch25h* homologs do not exhibit the same antiviral upregulation (25).

To further validate our findings in this zebrafish model, we evaluated CH25H expression in post-mortem human foetal brain samples with SARS-CoV-2-associated brain haemorrhages (9). First, we quantified microbleed number and size in the brain cortical samples, observing a variable size of microbleed which we classified in three area size categories (Fig 2A). Measuring the number and size of microbleeds, we classified the cortical samples by a bleeding score, where non-haemorrhagic samples had a score 0, and haemorrhagic samples were divided into scores 1 and 2. Score 2 samples had significantly more bleeds, quantified by total density number (Fig. 2B), and significantly more bleeds of medium size (Fig. 2C). We then evaluated CH25H levels by immunohistochemistry, observing a lack of CH25H+ cells in non-haemorrhagic samples and variable numbers of CH25H+ cells in haemorrhagic samples, both proximal and distal to haematomas (Fig. 2D). A significant increase in CH25H+ cells was observed in samples with a bleeding score of 2 (Fig. 2E). Altogether, our data shows that CH25H upregulation is part of an antiviral response in SARS-CoV-2-associated haemorrhages in developing zebrafish and human brain.

**Figure 2.**
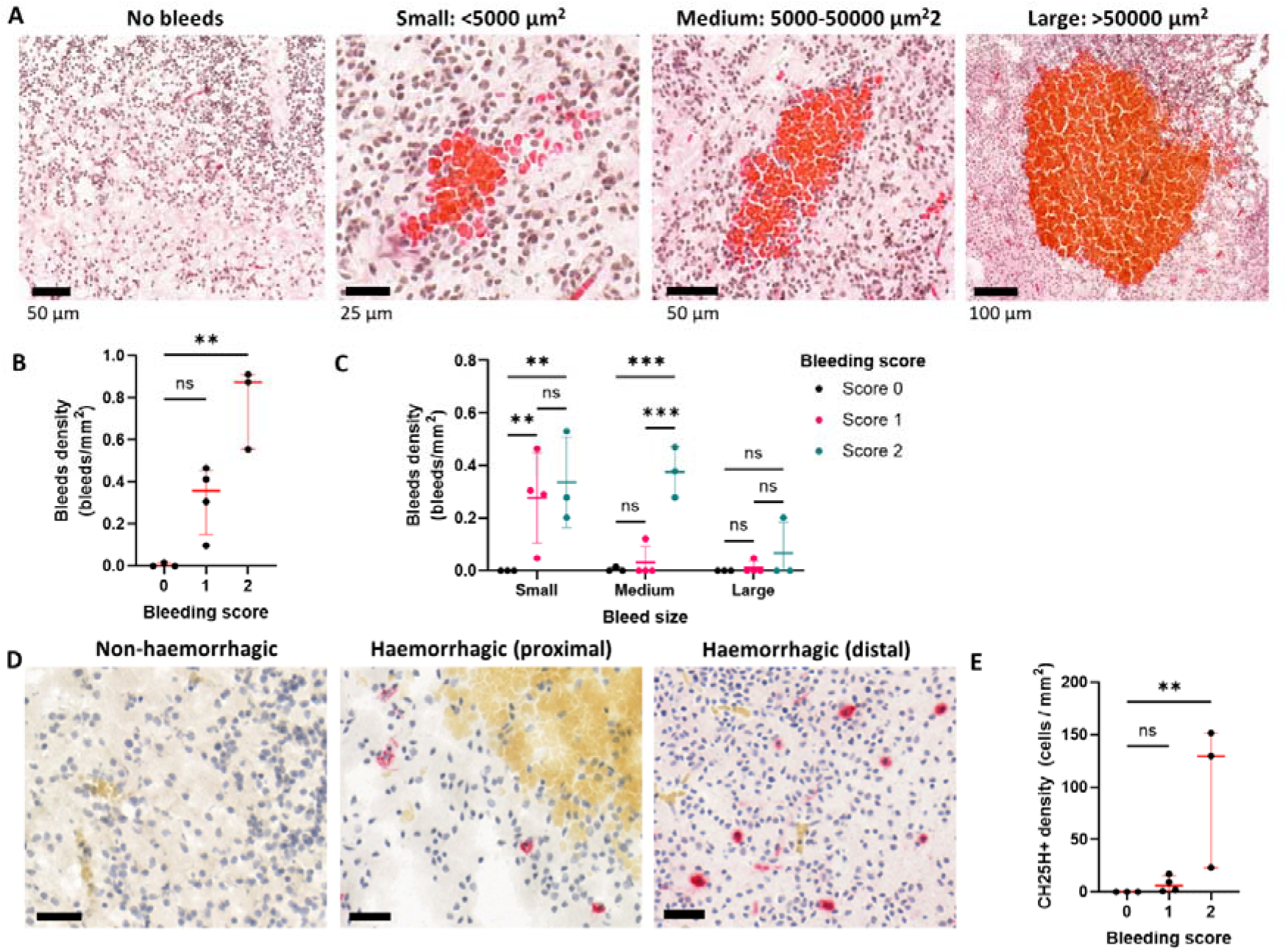
CH25H expression is associated with human foetal SARS-CoV-2-associated brain microbleeds. **A** Representative images of microbleeds found in human foetal cortex by H&E staining. Bleeds were classified into small, medium and large sizes according to area. Scale bar size is shown in each image. **B** Samples were categorised according to a bleeding score, based on bleed density. Control samples were classified in score 0, and haemorrhagic samples were classified in scores 1 and 2. **C** Density of small, medium, and large bleeds in samples with bleeding scores 0 to 2. **D** Representative images for CH25H staining (pink) and bleeds (yellow) from non-haemorrhagic or haemorrhagic samples, proximal and distal to bleeds. Scale bars 30 µm. **E** CH25H+ cell density was quantified in samples with bleeding scores 0 to 2. Data are mean ± S.E.M. **, p < 0.01; ***, p < 0.001; ***, p < 0.0001; ns, nonsignificant, determined by one-way ANOVA with Kruskal-Wallis post hoc test compared to score 0 (**B, E**), or two-way ANOVA with Tukey’s post hoc analysis multiple comparisons between scores (**C**).

### 25HC increases the severity of bleeding in a statin-dependent zebrafish ICH model

We next aimed to evaluate whether the CH25H metabolite 25HC could exacerbate cerebrovascular dysfunction. Inhibition of 3-hydroxy-3-methylglutaryl-CoA reductase (HMGCR) using statins, induces spontaneous brain haemorrhages in zebrafish larvae (13, 26, 27). We have previously shown that a reduced expression of *hmgcr* in a zebrafish model of type I interferonopathy leads to increased susceptibility to statin-induced ICH (13). As 25HC decreases expression of *HMGCR* by inhibition of the transcription factor SREBP2 (Sterol regulatory element-binding protein 2) (28), we hypothesised that 25HC would also exacerbate an ICH phenotype.

Wild type (WT) zebrafish larvae that were incubated with 25HC (25 µM, 24 h) exhibited significantly reduced transcription of the *HMGCR* zebrafish homolog gene *hmgcrb* (Fig. 3A). This confirmed that 25HC inhibited SREBP2 in zebrafish. To evaluate the effects of 25HC and ATV on brain bleeding in zebrafish, ATV-induced haemorrhages were identified by O-dianisidine staining (Fig. 3B) (13). Following ATV incubation, 25HC was injected (1 nl, 1 mM) directly into the bloodstream through the duct of Cuvier (29). 25HC injections alone did not increase ICH occurrence in untreated larvae, however, in ATV-incubated larvae we observed a non-significant increase in ICH frequency (Fig. 3C). To assess the extent of cerebral bleeding, we next evaluated brain haematoma area in the same animals, showing that in ATV treated embryos, 25HC induced a significant increase in bleed area (Fig. 3D). This zebrafish ICH model demonstrates that 25HC and ATV interact and exacerbate neurovascular instability.

**Figure 3.**
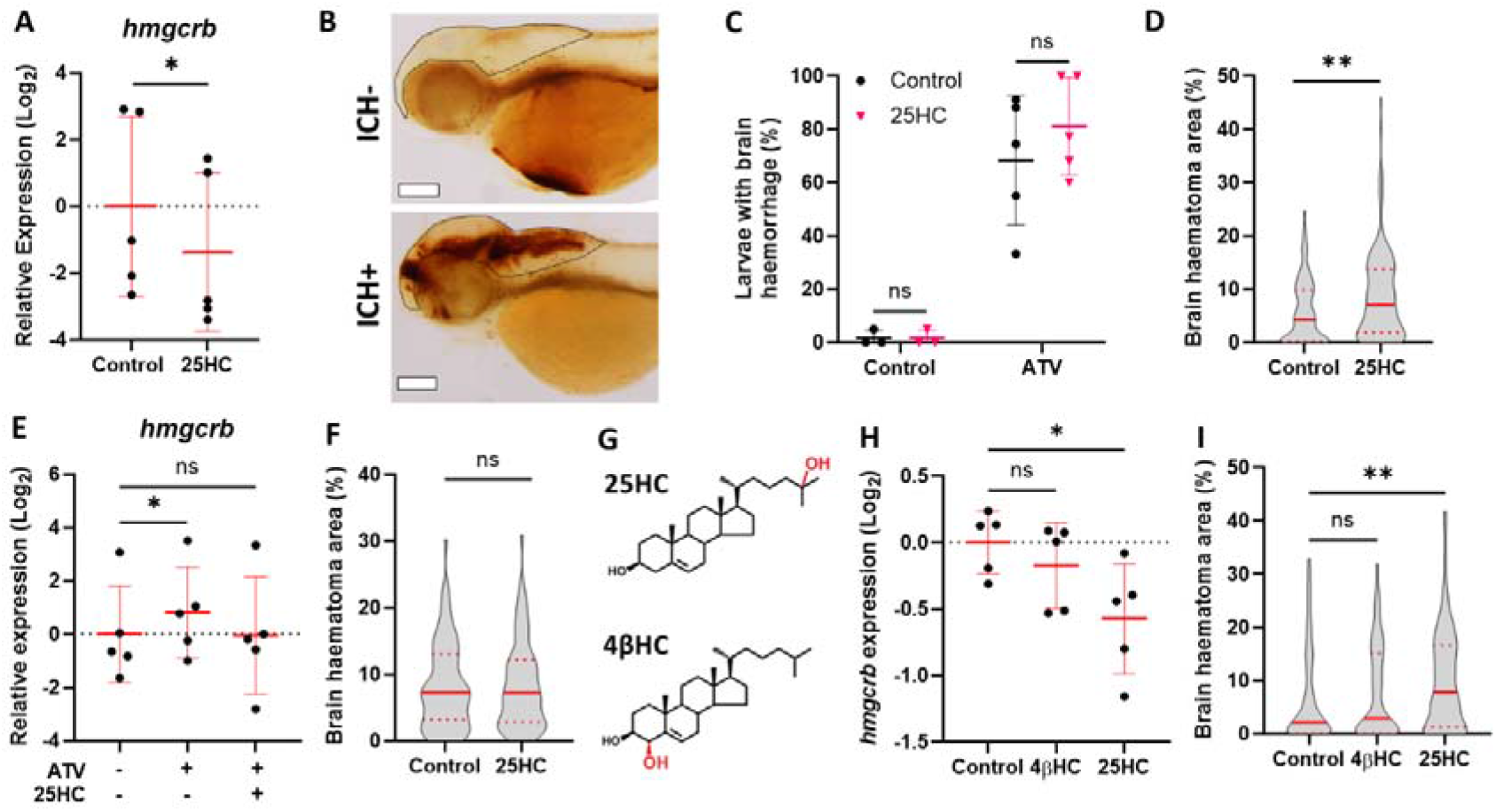
25HC worsens brain haemorrhage expansion in statin-induced ICH zebrafish model. **A** *hmgcrb* expression in 2 days post-fertilization (dpf) WT zebrafish larvae incubated with 25HC (25 μM, 24 h). **B-D** WT larvae were incubated in ATV (1 µM) at 28 hours post-fertilisation (hpf) and intravenously injected with 25HC (1 nl, 5 µM) at 32-36 hpf. Next day larvae were stained with O-dianisidine. Representative images of larvae without (ICH-) and with brain haemorrhage (ICH+) (**B**), ICH+ frequency per experiment (**C**) and brain haematoma area per larvae (**D**) were quantified (n=128 embryos, 5 independent experiments). Dotted lines indicate brain area. Scale bars 150 µm. **E** hmgcrb expression in 2 dpf WT larvae incubated with 25HC (25 µM) and ATV (1 µM) for 24 h. **F** *bbh* zebrafish larvae were injected with 25HC (1 nl, 5 µM) at 32-36 hpf. After 24 h larvae were stained with O-dianisidine and brain haematoma area was quantified (n=69-82 embryos, 3 independent experiments). **G** Comparison of 25HC and 4βHC structures. **H** Expression of *hmgcrb* in 2 dpf WT larvae incubated with 4βHC or 25HC (25 µM, 24h). **I** WT larvae were incubated in ATV (1 µM) at 28 hpf and injected with 4βHC or 25HC (1 nl, 2.5 µM) at 32-36 hpf. Next day larvae were stained with O-dianisidine and haematoma area was quantified (n=87-93 embryos, 3 independent experiments). Data expressed as mean⍰± ⍰SD (**A, C, E, H**) or median ± ⍰IQR (**D, F, I**). *, p < 0.05; **, p < 0.01; ns, nonsignificant, determined by paired t-test (**A**), randomized block two-way ANOVA with Sidak’s post-hoc test compared to control (**C**), Mann-Whitney test (**D, F**), randomized block one-way ANOVA with Dunnet’s post-hoc test compared to control (**E, H**), or Kruskal-Wallis test with Dunn’s post-hoc test compared to control (**I**).

We then evaluated whether the ATV and 25HC interaction was dependent on *HMGCR* inhibition. ATV incubation increased the expression of *hmgcrb*, a feedback response induced through the activation of SREBP2 (30), and 25HC co-incubation inhibited this process (Fig. 3E). This suggested that 25HC increased the sensitivity to ATV by decreasing *HMGCR* levels. To confirm whether the 25HC phenotype was dependent on ATV inhibition of *HMGCR*, we repeated the experiment in an alternative ICH zebrafish model. The homozygous *bbh* zebrafish mutant expresses a hypomorphic mutation in the *βpix* gene which leads to dysfunctional neuroendothelium, resulting in a comparable ICH phenotype to the ATV model independently of *HMGCR* activity (31). 25HC injections in *bbh* larvae caused no differences in haematoma size (Fig. 3F). This demonstrated that 25HC effects on neurovascular stability are seemingly dependent on *HMGCR* inhibition. To confirm this, we compared 25HC to another oxysterol, 4β-hydroxycholesterol (4βHC), which shares a similar structure (Fig. 3G) but has no effect on SREBP2 activity (17). This was corroborated by *hmgcrb* expression in WT zebrafish larvae, which was significantly inhibited by 25HC incubation but not by 4βHC (Fig. 3H). When both oxysterols were injected alongside ATV incubation, only 25HC increased haematoma area in WT zebrafish larvae (Fig. 3I). These results revealed that 25HC increases the risk of ATV-induced ICH by inhibiting SREBP2 and reducing *hmgcrb* expression.

### 25HC remodels cholesterol metabolism in human brain endothelial cells

To further explore the effects of 25HC in brain ECs, we used the human brain microvasculature cell line hCMEC/D3. Our zebrafish data (Fig. 3) suggested that 25HC effects were dependent on SREBP2 inhibition. As SREBP2 is a master regulator of cholesterol biosynthesis, we evaluated the expression of several cholesterol synthesis enzymes, including *HMGCR*, in hCMEC/D3 cells at 4 and 24 hours after 25HC exposure (Fig. 4A). The expression of the enzymes *CYP51A1* and *EBP* was significantly decreased in hCMEC/D3 cells after 24 hours exposure, whilst the expression of *HMGCR* and *SQLE* was significantly decreased after both 4 and 24 hours exposure (Fig. 4A). 25HC also regulates cholesterol metabolism through a second transcription mechanism, activating LXR transcription factors (17, 28). ABCG1 is an LXR target involved in cholesterol efflux (33). Using the same samples (Fig. 4A), we detected significant upregulation of ABCG1 expression after exposure to 25HC for both 4 and 24 hours (Fig. 4B).

**Figure 4.**
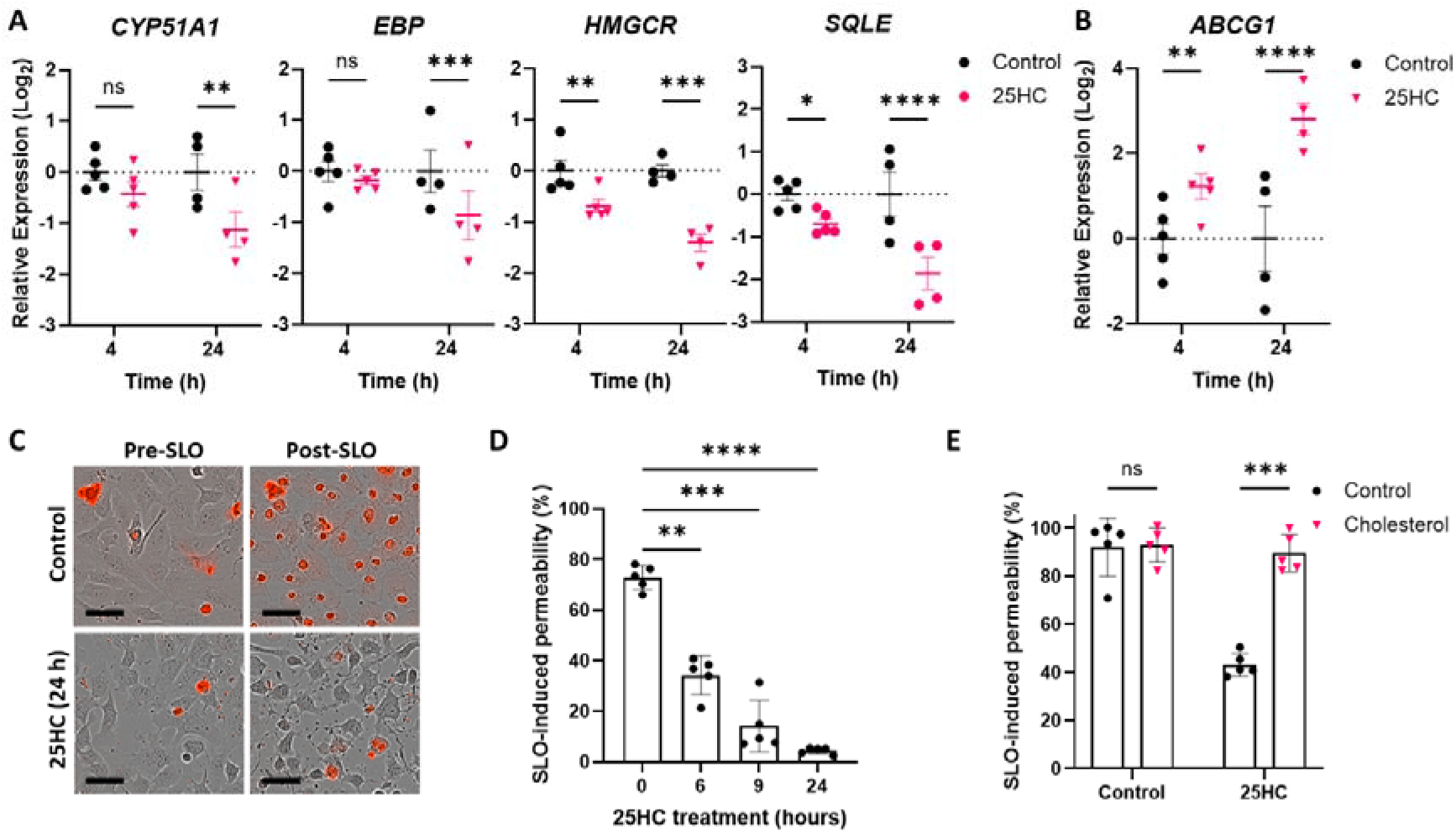
25HC remodels cholesterol metabolism in human brain endothelial cells. **A-B** Gene expression of *HMGCR, SQLE, CYP51A1, EBP* (**A**), and *ABCG1* (**B**) in hCMEC/D3 cells after 25HC treatment (5 μM, 4 and 24 h). **C-D** hCMEC/D3 cells were pre-treated with 25HC (5 μM, 0 to 24 h) before incubation with streptolysin O (SLO). Membrane permeability was measured by ToPro3+ uptake (red signal) before and after SLO, representative images (**C**) and quantification (**D**) are shown. Scale bars 37.5 µm. **E** Permeability of hCMEC/D3 cells, pre-treated with 25HC (5 μM, 6 h) and then with soluble cholesterol (80 μM, 1 h), after streptolysin O (SLO) incubation. Data expressed as mean⍰± ⍰SD. *, p < 0.05; **, p < 0.01; ***, p < 0.001; ****, p < 0.0001; ns, nonsignificant, determined by paired t-test (**A, B**), randomized block one-way ANOVA with Dunnet’s post hoc test compared to 0 µM (**D**), or randomized block two-way ANOVA with Sidak’s post-hoc test compared to control (**E**).

Besides transcriptional regulation, another important antimicrobial mechanism induced by 25HC is the depletion of plasma membrane-accessible cholesterol levels (17, 18, 32). Using Streptolysin O (SLO), a microbial toxin that depends on accessible cholesterol to form membrane pores (32), we indirectly evaluated accessible cholesterol levels by measuring SLO-induced permeability through uptake of the small dye ToPro3 (Fig. 4C). SLO permeabilised the membrane of hCMEC/D3 cells, and this permeability was inhibited by 25HC pre-treatment in a time-dependent manner (Fig. 4C-D). This suggested that 25HC quickly induced the depletion of accessible cholesterol. To confirm this, cells were supplemented with soluble cholesterol (80 µM, 1h) after 25HC treatment. Cholesterol supplementation rescued the permeability phenotype (Fig. 4E), thereby confirming that the reduction in permeability was due to a decrease in accessible cholesterol levels. Altogether, these results suggest that 25HC not only remodels cholesterol metabolism of brain ECs by targeting *HMGCR* but also by inhibiting cholesterol synthesis, modulating cholesterol efflux and decreasing accessible cholesterol levels.

### 25HC-induced dysfunction of human brain endothelial cells depends on cholesterol metabolism

As 25HC exacerbated the neurovascular instability phenotype in zebrafish (Fig. 3), we next questioned whether 25HC would also affect the barrier function of brain ECs *in vitro*. For this, we analysed the permeability of fluorescein dextran 70 kDa (FD70), a method previously used to assess hCMEC/D3 barrier function (33). 25HC induced a concentration-dependent decrease of barrier function, with 25HC (5 µM) treatment leading to a significant 85% increase in FD70 permeability (Fig. 5A). To further investigate the impact of 25HC on endothelial function, we used a previously established scratch assay to evaluated hCMEC/D3 cell migration (34) (Fig. 5B). One day pre-treatment with 25HC decreased cell migration in a concentration-dependent manner (Fig. 5C). These effects were neither associated to cell death (Fig. s1A-B) not to a decrease in cell density, as 25HC treatment did not affect total cell numbers (Fig. s1C) and inhibition of cell migration in the scratch assay persisted even with cell cycle arrest induced by mitomycin C (Fig. s1D).

**Figure 5.**
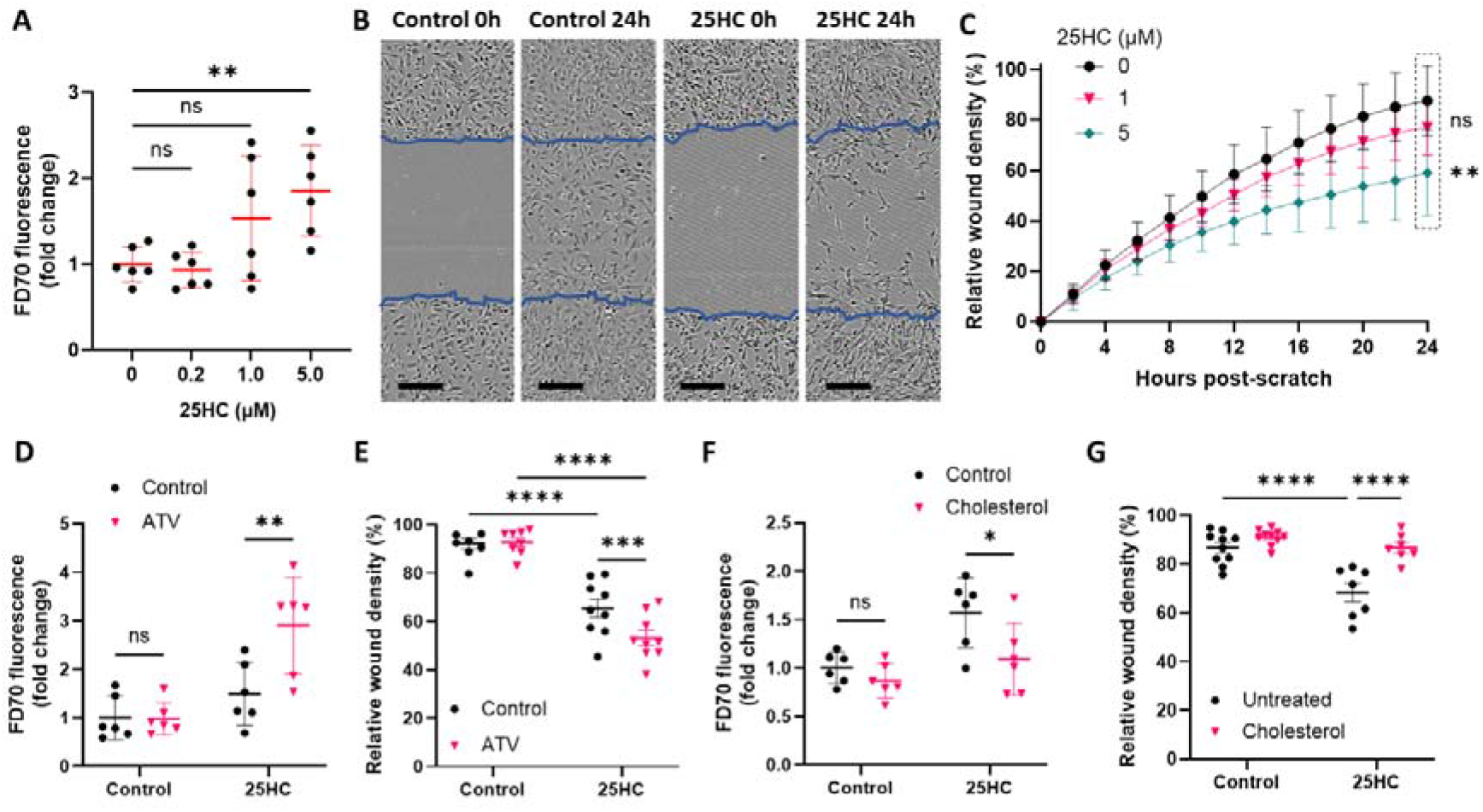
25HC dysregulates the function of human brain endothelial cells. **A** Permeability of fluorescein-conjugated dextran 70 kDa (FD70) in hCMEC/D3 cell monolayer pre-treated with different concentrations of 25HC (0-5 μM, 24h). **B-C** hCMEC/D3 cell migration in a scratch assay, cells were pre-treated with different concentrations of 25HC (0-5 μM, 24 h before scratch). Representative images are shown for cells 0 and 24 hours after scratch (**B**, 25HC 5 μM). Blue lines show initial scratched area. Scale bars 100 µM. Migration was quantified as relative wound density (**C**, n=10). **D** FD70 permeability was analysed in hCMEC/D3 cells treated with 25HC (5 μM, 24 h) and ATV (1 μM, 24 h). **E** Cell migration at 24-hours post-scratch of hCMEC/D3 cells pre-treated with 25HC (5 μM, 24h) before scratch and treated with ATV (1 µM, 24h) after scratch. **F** Fluorescein-conjugated dextran 70 kDa (FD70) permeability was analysed in hCMEC/D3 cells pre-treated with 25HC (5 μM, 14h) and then supplemented with soluble cholesterol (80 µM, 1h). **C** Cell migration at 24-hours post-scratch of hCMEC/D3 cells pre-treated with 25HC (5 μM, 24 h) and then supplemented with cholesterol (80 µM, 2 h) before scratch. Data expressed as mean⍰± ⍰SD. *, p < 0.05; **, p < 0.01; ***, p < 0.001; ns, nonsignificant, determined by randomized block one-way ANOVA with Dunnet’s post-hoc test compared to 0 µM (**A**), matched measures two-way ANOVA with Dunnet’s post-hoc test compared to 0 µM (**C**), or randomized block two-way ANOVA with Sidak’s post-hoc test compared to control (**D**-**G**).

We next aimed to determine if HMGCR inhibition was involved in the 25HC-induced dysfunction of hCMEC/D3 cells. As in the zebrafish ICH model, HMGCR inhibition by statins has been shown to decrease *in vitro* EC barrier function at micromolar concentrations (27, 35). Therefore, we hypothesised that 25HC and ATV would also interact in hCMEC/D3 cells. ATV treatment alone (1 µM, 24 h) did not affect barrier function, while ATV and 25HC co-treatment significantly increased permeability in comparison to 25HC alone (Fig. 5D). A similar interaction between 25HC and ATV was observed in the scratch assay. ATV alone had no effect on cell migration, while it significantly exacerbated inhibition of cell migration in 25HC pre-treated cells 24 hours post-scratch (Fig. 5E). These results demonstrate that similar to our zebrafish model, 25HC-induced hCMEC/D3 dysfunction is dependent on *HMGCR* inhibition.

As our *in vitro* analysis suggested that 25HC also decreases cholesterol synthesis and cholesterol accessible levels in brain ECs, we also aimed to determine whether 25HC effects on endothelial function would depend on cellular cholesterol. For this, hCMEC/D3 cells were supplemented with soluble cholesterol (as in Fig. 4E) after being pre-treated with 25HC. Using the FD70 permeability assay, we observed that cholesterol supplementation rescued the effect of 25HC on barrier function (Fig. 5F). Similarly, cholesterol supplementation also rescued 25HC inhibition of cell migration. hCMEC-D3 cells were pretreated with 25HC for 1 day and then supplemented with cholesterol before the scratch. This treatment significantly increased the migration of 25HC-treated cells (Fig. 5G). Altogether, these results suggest that the 25HC-induced dysfunction of brain ECs is also mediated by a decrease in cholesterol availability.

## Discussion

Here, we propose that the CH25H/25HC pathway represents an important component of brain EC dysfunction associated with infection. Our findings in zebrafish larvae and human foetal samples suggest that CH25H is upregulated in viral-associated developmental brain haemorrhages. In these samples, most human foetal haemorrhages occurred between 12 and 14 weeks post-conception (9), a critical window for neurovascular development marked by endothelial junction protein upregulation (49, 50). Similarly, in our zebrafish ICH models, bleeding occurs between 2-3 dpf (23, 26, 31), coinciding with brain angiogenesis and barrier function development (51). These suggest that an antiviral response, including upregulation of CH25H, may affect critical steps of brain vascular development.

We show that the CH25H metabolite, 25HC, can induce brain EC dysfunction *in vitro* and exacerbates ICH in a zebrafish model. These findings align with previous research linking the CH25H/25HC pathway to vascular inflammation seen in atherosclerosis, EAE, and lung injury (19-21). While these previous studies have primarily associated the CH25H/25HC pathway to ER stress (21) or regulation of inflammatory pathways (19, 20), our data highlight cholesterol remodelling as a key process controlling brain endothelial function. 25HC modulated cholesterol metabolism at a transcriptional level and by reducing accessible cholesterol levels in the plasma membrane. Moreover, 25HC-induced brain EC dysfunction was linked to *HMGCR* activity and cholesterol availability.

Our findings are consistent with previous reports highlighting the importance of cholesterol homeostasis for EC function, as dysregulation of cholesterol synthesis or transport has been shown to compromise angiogenesis and barrier function (36-38). Endothelial dysfunction has been previously associated with cholesterol-dependent defects in adherence junction proteins (36, 39), but 25HC may also disrupt neurovascular responses to blood flow, as cholesterol in the plasma membrane is crucial for endothelial signalling in shear stress responses (40, 41). Cholesterol remodelling could also affect other processes dependent on intermediate metabolites, such as protein prenylation, which depends on isoprenoids produced by HMGCR (42). The vascular instability induced by ATV in our *in vitro* and zebrafish models has been linked to prenylation defects in endothelial GTPases (26, 35, 43). It is possible that the interaction between ATV and 25HC could also have been mediated by defective prenylation in our models, as decreased prenylation has also been proposed as a potential antiviral mechanism for 25HC (44).

While in this study we used exogenous 25HC in zebrafish and *in vitro* models to study the effects of cholesterol remodelling on brain endothelial function, future investigations should assess the role of CH25H upregulation and endogenous 25HC production in neurovascular function. In response to inflammation, CH25H is upregulated and 25HC levels increase within the brain, as observed in murine models involving lipopolysaccharide (LPS) challenge (45) and tauopathy neurodegeneration (46). CH25H is upregulated in microglia (46, 47), and secretion of 25HC from LPS-stimulated microglia has been observed *in vitro*. Future studies should explore whether microglia-derived 25HC contributes to neurovascular function, similar to effects reported in atherosclerosis where macrophage-derived 25HC regulates vascular inflammation (19). Alternatively, intrinsic endothelial CH25H expression could also regulate neurovascular function, as CH25H upregulation has been detected in brain ECs in response to brain inflammation (20, 48) and endothelial CH25H has been shown to modulate EAE progression (20).

While our study used foetal human brain samples and zebrafish larval ICH models, the mechanisms involving viral infection as a risk factor of adult ICH may differ from those in the developing brain. Animal models of infection-triggered brain haemorrhages are required to understand this process. Although brain haemorrhages in adult mice have been observed following Japanese encephalitis and dengue infections (52, 53), these are associated with strong encephalitis and don’t resemble milder ICH cases triggered by flu-like infections (8). Mild viral infection may only trigger adult ICH in combination with other risk factors and/or co-morbidities. Interestingly, hypocholesterolaemia is a known ICH risk factor (54-57), and statins have long been debated as potential contributors to ICH risk (58-61). Since our results suggest that the effects of 25HC in brain ECs are dependent on cholesterol remodelling, it is possible that infection-induced upregulation of the CH25H/25HC pathway could interact with hypocholesterolaemia and statin use in the development of adult-onset ICH.

In conclusion, we have reported the upregulation of CH25H in viral-associated developmental brain haemorrhages in a zebrafish model and in human foetal brain samples. Using *in vitro* and zebrafish models, we have demonstrated that 25HC dysregulates brain EC function by remodelling cellular cholesterol. Our results highlight a novel association between 25HC, cholesterol remodelling and brain EC dysfunction.

## Material and Methods

### Zebrafish husbandry

Zebrafish were raised and maintained at The University of Manchester Biological Services Unit under standard conditions as previously described (62). Adult zebrafish husbandry was approved by The University of Manchester Animal Welfare and Ethical Review Board. All experiments were performed per U.K Home Office regulations (PPL: PP1968962). The zebrafish used in this study were raised and maintained at The University of Manchester Biological Services Unit, under standard conditions, with adults housed in mixed-sex tanks with a recirculating water supply maintained at 28°C under a 14/10 hour light/dark cycle, as previously described (62). Wild type AB, double transgenic Tg(fli1a:GFP)^y1^ / (gata1a:DsRed)^sd2^ (63, 64) and βpix mutant *bbh* (*bbh*^m292^) (31) adult zebrafish lines were used. Fertilized eggs were collected following natural spawning and incubated at 28 °C in fresh E3 medium, except embryos injected with SARS-CoV-2 spike protein which were kept at 26 °C until 48 hours post-fertilization (hpf) to delay development. The embryos were staged according to standard guidelines (65). Larvae were anaesthetized with Tricaine Methanesulfonate (MS222) 0.02% before injections and imaging. After termination of the experiment, all embryos were killed before protected status (5 dpf) using a lethal dose MS222 (4%) and freezing at -20 °C.

### Zebrafish larvae ICH models

For the SARS-CoV-2 spike 1 protein model, Tg(fli1a:GFP)^y1^ / (gata1a:DsRed)^sd2^ larvae were kept at 26°C, dechorionated at 24 hpf and incubated with 0.3% N-phenylthiourea (PTU; Sigma-Aldrich) to inhibit melanogenesis. At 48 hpf, larvae were injected in the hindbrain with 2 nl of SARS-CoV-2 spike S1 protein (0.25 mg/ml) (SinoBiologicals, 40591-V08H3, molecular weight of 116 kDa) or BSA (0.43 mg/ml) (Sigma-Aldrich, molecular weight of 66.4 kDa) in PBS supplemented with phenol red 0.05%. Following hindbrain injections, larvae were kept at 28 °C for 24 h and brain haemorrhages were imaged *in vivo*.

For the 25HC and ATV model, zebrafish WT larvae were dechorionated and treated by immersion with ATV 1 µM (Merck) at 28 hpf. Control groups included vehicle hydroxypropyl-β-cyclodextrin (HβCD) in all experiments. ATV-treated WT or *bbh* embryos at 32 to 36 hpf were injected with 1 nl of 25HC or 4βHC (0.5 or 1 mM), HβCD 2.25%, phenol red 0.05% in PBS through the Duct of Cuvier (29). To visualize bleeds, embryos were stained at 54 hpf using an o-dianisidine (Sigma) protocol, as previously described (26). Stained embryos were mounted in 50% glycerol PBS and imaged. For qPCR analysis, embryos were treated by immersion with 25HC or 4βHC 25 µM, alongside 0.11% HβCD, at the same time as ATV.

Imaging was performed in a Leica M165FC light stereo microscope with DFC7000T camera and processed using LASX software (version 3.3.3.16958). Images were analysed using ImageJ, brain area (excluding the eyes) was manually selected and threshold tool was used identically on all conditions to quantify brain haemorrhage area. Injections and manual selection of brain area were conducted blind to treatment.

### Human foetal cortical samples

Human foetal tissues were obtained from the Human Development Biology Resource (HDBR), provided by the Joint MRC/Wellcome Trust (grant #MR/R006237/1) (www.hdbr.org). The HDBR provided fresh tissue from fetuses aged 9-21 post-conception weeks (pcw). Brains were fixed for at least 24 hours at 4°C in 4% (wt/vol) paraformaldehyde (PFA) in 120 mM phosphate buffer (pH 7.4). Brain cortexes were then sucrose-treated (15 and 30% sucrose solution sequentially for 24 h each), OCT-embedded, and then 20 μm thick sections were cut using a cryostat, as previously described (9).

### Staining of human foetal samples

To visualise microbleeds, slides were stained with a haematoxylin and eosin (H&E) standard protocol and mounted with DPX mounting medium. For CH25H immunohistochemistry, antigen retrieval was performed by placing slides in 97.5 °C water bath in Tris-EDTA (pH 9.0) for 20 min. Non-specific binding was blocked with PBS with 5% goat serum, 0.3% Triton X-100 and 0.1% Tween for 1 h. CH25H antibody (Aviva system biology, OABF01697) was diluted at 1:2000 blocking solution and incubated on slide overnight at 4 °C. Slides then were incubated with biotinylated goat anti-rabit secondary antibody (BA-1000, Vector Laboratories) diluted in TBST buffer (tris-buffered saline with 0.1% tween and 0.1% BSA) at 1:400 for 90 min at RT. Slides then were stained with alkaline phosphatase ABC-AP Kit (AK-5000, Vector Laboratories), following manufacturer instructions. Finally, slides were counterstained with haematoxylin and mounted with DPX. Slides were imaged using a 3D Histech Pannoramic 250 Flash Slide Scanner. Images were analysed using the software QuPath (version 0.5.0). Bleed and total brain areas were manually selected, and CH25H+ cells were selected using a custom script.

### Cell culture

The immortalised human cerebral microvascular endothelial cell line hCMEC/D3 cells (Merck) were cultured in endothelial cell growth medium MV (Promocell, C-22020) and PenStrep (100 units/ml penicillin and 100 μg/ml streptomycin) at 37 °C in a humidified atmosphere containing 5% CO2. Flask, plates and well inserts were pre-coated with rat tail collagen type I (Merck, UK), (1:100 in PBS) at 37 °C for 1 h, before hCMEC/D3 seeding. Cells were passaged at 100% confluency using Trypsin-EDTA solution (Sigma) and were used in experiments until passage 15.

hCMEC/D3 cells were seeded at a density of 71,000 cells/cm^2^ overnight in all assays, with the exception of permeability and scratch assays (specific cell density stated in their subsections). Cells then were treated with 25HC (Sigma-Aldrich, H1015), soluble cholesterol (Sigma-Aldrich, C4951), Staurosporine (Cell guidance system, SM97), Mitomycin C (Roche, 10107409001) or ATV (Merck, SML3030), at concentrations and times stated in figure legends. When used as a carrier, control groups included ethanol or DMSO at the same concentration as treated groups.

### Quantitative PCR

Total RNA was pooled groups of zebrafish larvae (n=30 larval heads, n=15 full larvae) or extracted from hCMEC/D3 cells seeded in 12 well/plates using a standard TRIzol (Invitrogen) method. Complementary DNA (cDNA) was synthesized from 800ng RNA as previously described (18). Quantitative PCR (qPCR) was performed on a StepOne Plus Real Time PCR machine (Applied Biosystems). cDNA samples were analysed using power SYBR green mastermix (Applied Biosystems) and primers (Thermo Fischer). Gene expression was normalized by geometric averaging of two internal control genes. Reference genes used were *hprt1* and *actb2* for zebrafish samples, and *HPRT1* and 18S for human samples. A Taqman (Applied Biosystems) protocol and probes were used for hmgcrb and sqlea zebrafish genes, using *hprt1* as a reference gene. List of primers and Taqman probes in supplementary table 1.

### Streptolysin O (SLO) assay

hCMEC/D3 cells were seeded in 96 well/plates overnight and treated as previously stated. After cell treatments, media was changed to Opti-MEM reduced serum media (Thermo Fisher, 11058021) with To-Pro-3 0.5 µM (Thermo Fisher, T3605). SLO (Sigma-Aldrich, SAE0089) was activated with DTT 20 mM (30 min, 37 °C) before adding it to cells at 2 U/µl. Images were captured before SLO addition, and every 15 min after SLO addition using an IncuCyte ZOOM System (Essen Bioscience) with a ×20/0.61 S Plan Fluor objective. 90 min after SLO addition, lysis solution (Promega, G1780) was added to capture an image with 100% permeability. Images were automatically analyzed using the IncuCyte software (Essenbio), using custom scripts which gave optimal detection for To-Pro-3 positive cells that ran identically on all conditions.

### Permeability assay

hCMEC/D3 cells were seeded overnight at 50,000 cells/well in CellQart 24-well inserts with 0.4 µM pore size (Sterlitech, 9320412) and treated as previously stated. After cell treatments, media was changed to Opti-MEM reduced serum media, adding fluorescein dextran 70 kDa (FD70, 0.1 mg/ml) (Sigma-Aldrich, 46945) inside CellQart insert. 30 min later, fluorescence of media outside CellQart insert was measured using a Fluorstar reader (BMG Labtech).

### Scratch assay

hCMEC/D3 cells were seeded in 6 well/plates overnight and pre-treated with 25HC for 24 h before re-seeding for scratch assay. hCMEC/D3 cells were seeded at 40,000 cells/well in 96 well ImageLock plates (Essen BioScience) and left to adhere for 4 h before scratch. During that time, cells were pre-treated with 25HC, mitomycin 5 µg/mL or soluble collagen for 1 or 2 h before scratch. Scratch wound injury was carried out using a 96-pin IncuCyte WoundMaker Tool (Essen BioScience). Cells were then washed twice with PBS and replaced with media. Phase contrast images were acquired at 2 h intervals for a period of 24 h using an Incucyte Zoom Live Cell Analysis system with a 4×/3.05 Plan Apo OFN25 objective. The 96-well Cell Migration Software Application Module (Essen BioScience) was used to quantify relative wound density, as previously described (34).

### Western blot

Western blot analysis was performed on hCMEC/D3 cell lysates using antibodies against full-length and cleaved caspase-3 (Abcam, ab32351, 1:500 dilution), VE-Cadherin (Cell Signalling, D87F2, 1:500 dilution) and β-Actin (Sigma-Aldrich, A3854, 1:10,000 dilution). Samples were run on 8% (Ve-Cadherin) or 12% (Caspase-3) SDS-polyacrylamide gels. Gels were transferred using a Trans-Blot® TurboTM Transfer System (Bio-Rad) before blocking with 5% BSA in PBST (PBS, 1% Tween 20) for 1 h at room temperature. Membranes were washed and incubated (4 °C) overnight in primary antibody in PBST with 0.1% BSA. Following this, membranes were washed and incubated with horseradish peroxidase–conjugated secondary antibodies (Dako) in PBST with 0.1% BSA for 1 h at room temperature. Finally, membranes were washed, incubated in ECL Western Blotting Detection Reagent (GE Life Sciences) and imaged using a G:BOX gel doc system (Syngene).

### Statistical analysis and data presentation

Statistical analysis was performed using GraphPad Prism (v10.2). Data were presented as single data points with mean⍰±⍰SD, except for violin plots with median and ±⍰IQR. Experimental replicates (n) were defined as experiments performed on embryo clutches produced by different zebrafish adult pairs, individual zebrafish embryos, individual human donors, or different hCMEC/D3 passages. Experimental replicates were matched when measured several times (repeated measures ANOVA) or obtained from the same embryo clutches or hCMEC/D3 passages (randomized block ANOVA or paired t-test). Data were assessed for normal distribution using Shapiro-Wilk normality test. Parametric data were analysed using t-test or one-way ANOVA with Dunnett’s post-hoc test. Non-parametric data were analysed using Mann-Whitney test or Kruskal-Wallis test with Dunn’s post-hoc test. Two-factor data were analysed by two-way ANOVA with Sidak’s post-hoc test or with Tukey’s post-hoc test, time course data were analysed by two-way ANOVA with Dunnett’s post-hoc test.

## Acknowledgements

We thank the aquatics staff at the Biological Services Unit at The University of Manchester (UoM) for assistance with zebrafish care. We also thank the Bioimaging Core Facility at UoM.

## Funding

P.K. and V.S.T. were funded by Medical Research Council (MRC) grant MR/T03291X/1. C.B.L. and V.S.T. were funded by MRC grant MR/Y004183/1. S.E.W was funded by MRC Doctoral Training Programme studentship (MR/N013751/1). R.Z was funded by University of Manchester-China Scholarship Council Joint Scholarship. A.B. was funded by a 4 year British Heart Foundation PhD award (FS/4yPhD/F/22/34179).

## Conflict of Interests

The authors declare that there is no conflict of interest.

## Research ethics and patient consent

Human foetal tissues were obtained from the Human Development Biology Resource (HDBR), provided by the Joint MRC/Wellcome Trust (grant #MR/R006237/1).

## Data availability

The data of this study are available from the corresponding author upon reasonable request.

## Supplementary Files

**Table S1**. qPCR primers and probes

## Notes

### Competing Interest Statement

The authors have declared no competing interest.

### Summary of Updates

Manuscript title, abstract and main text have been revised to focus the analysis on mechanisms of cholesterol remodelling. The figures were revised, including changes in figure order, figure merging, and some figures being moved to supplementary data. Data was removed from the manuscript: SARS-CoV-2 spike analysis in human foetal samples (Figure 2) and VE-cadherin analysis in hCMEC/D3 cells (previous Figure 4). qPCR data was added in new figure 4.

