## Supplementary material for "25-hydroxycholesterol promotes brain endothelial dysfunction by remodelling cholesterol metabolism": Fig. s1

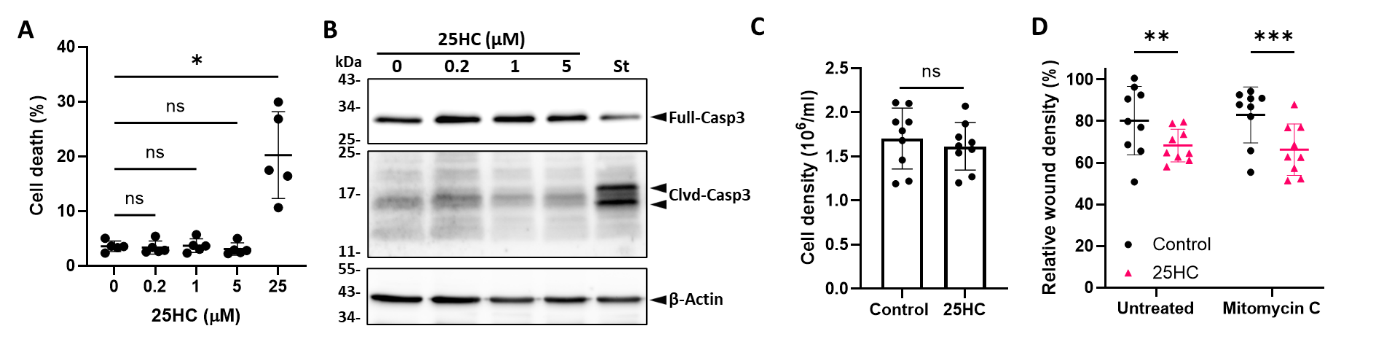

**Figure S1. 25HC remodels cholesterol metabolism in human brain endothelial cells. A** hCMEC/D3 cells were pre-treated with 25HC (0 - 25 μM) for 24 h, and cell death was assessed by loss of membrane integrity, measuring ToPro3+ uptake. **B** Western blot analysis of hCMEC/D3 cell lysates pre-treated with 25HC (0-5 μM, 24 h), compared to staurosporine 5 µM (St) as apoptosis-positive control. Blots for full caspase-3 (FL-Casp3), activated cleaved caspase-3 (Clvd-Casp3) and β-actin as loading control, image is representative of three independent experiments. **C** hCMEC/D3 cell density after treatment with 25HC (5 μM, 24 h), counted using a haematocytometer after trypsin detachment. **D** hCMEC/D3 cell migration 24 hours post-scratch, pre-treated with 25HC (5 μM, 24 h before scratch) and then mitomycin D (5 µg/ml, 2 h before scratch).
