## supplementary table 1 for "25-hydroxycholesterol promotes brain endothelial dysfunction by remodelling cholesterol metabolism"

**Table S1**. qPCR primers and probes

| Zebrafish primers | | Sequence |
| --- | --- | --- |
| *hrpt1* | forward | TTGCAGTAGCTTGTCGGTGT |
| *hrpt1* | reverse | CAGACGTTCAGTTCGGTCCA |
| *actb2* | forward | ATGGATGATGAAATTGCCGCAC |
| *actb2* | reverse | ACCATCACCAGAGTCCATCACG |
| *ch25h* | forward | CGGTGAATCCCATGTTGCTT |
| *ch25h* | reverse | AGCTCCTCCGTAAAGTCCAAAA |
| *ch25hl1.1* | forward | GTACTGCTGGCCTTCTCCAG |
| *ch25hl1.1* | reverse | GGCATAGGCATTACCACGTT |
| *ch25hl1.2* | forward | CGACTCAACCACTCAGAGACC |
| *ch25hl1.2* | reverse | AAGGTACGGCAGGACAAGAA |
| *ch25hl2* | forward | CAATGTACCTGGTGCTGGTG |
| *ch25hl2* | reverse | GCAGATGGTTGTACGTGGTG |
| *ch25hl3* | forward | TCTTCTCGGTGCCCTTCTTA |
| *ch25hl3* | reverse | AAACGCCAACCACGTATTTC |
| **human primers** | | **Sequence** |
| *HPRT1* | forward | CAGGCGAACCTCTCGGCTTT |
| *HPRT1* | reverse | GGGTCGCCATAACGGAGCC |
| *18S* | forward | GTAACCCGTTGAACCCCATT |
| *18S* | reverse | CCATCCAATCGGTAGTAGCG |
| *HMGCR* | forward | GACGTGAACCTATGCTGGTCAG |
| *HMGCR* | reverse | GGTATCTGTTTCAGCCACTAAGG |
| *SQLE* | forward | CTCCAAGTTCAGGAAAAGCCTGG |
| *SQLE* | reverse | GAGAACTGGACTCGGGTTAGCT |
| *ABCG1* | forward | GAGGGATTTGGGTCTGAACTGC |
| *ABCG1* | reverse | TCTCACCAGCCGACTGTTCTGA |
| **zebrafish probes** | | **Assay ID** |
| *hprt1* | | Dr03095135_m1 |
| *hmgcrb* | | Dr03128326_m1 |
| *sqlea* | | Dr03131215_g1 |
